## supplemental data for "SRSF2 is a key player in orchestrating the directional migration and differentiation of MyoD progenitors during skeletal muscle development"

**Table S1. Antibodies used in IF, WB and ChIP analasis**

| | | | |
|---|---|---|---|
| Myosin heavy chain (MHC) Antibody | DSHB | Cat# mf-20-s | IF |
| RFP Antibody | Rockland | Cat# 600-401-379 | IF |
| Phalloidin | YEASEN | Cat# 40762ES75 | IF |
| Laminin-α2 Antibody | ENZO | Cat# ALX-804-190 | IF |
| Desmin Antibody | Santa Cruz | Cat# sc14026 | IF |
| Myog Antibody | Santa Cruz | Cat#(sc-576) | IF |
| Goat anti-Mouse IgG (H+L) Cross-Adsorbed Secondary antibody, Alexa Fluor 488 | Thermo Fisher Scientific | Cat# A-11001 | IF |
| Goat anti- Rabbit IgG (H+L) Cross-Adsorbed Secondary antibody, Alexa Fluor 546 | Thermo Fisher Scientific | Cat# A-11010 | IF |
| Alexa Fluor® 647-conjugated AffiniPure Donkey Anti-Rat IgG (H+L) | Jackson | Cat# 712-605-150 | IF |
| SRSF2 | Millipore | Cat# 04-1550 | WB |
| Aurka Antibody | Abcam | Cat# ab13824 | WB |
| CDK1 Antibody | Beyotime | Cat# AF1516 | WB |
| pRB Antibody | Beyotime | Cat# AF1564 | WB |
| p-pRB Antibody | Beyotime | Cat# AF1135 | WB |
| p27 Antibody | Cell Signaling Technology | Cat# 3698 | WB |
| AKT Antibody | Cell Signaling Technology | Cat# 4691 | WB |
| p-AKT Antibody | Cell Signaling Technology | Cat# 4060 | WB |
| p62 Antibody | Sigma-Aldrich | Cat# P0067 | WB |
| β-Actin Antibody | Santa Cruz | Cat# sc47778 | WB |
| Anti-HA tag | Abcam | Cat# ab9110 | ChIP |

**Table S2. Primer sequences used for RT-PCR**

| Gene | Forward (5’–3’) | Reverse (5’–3’) |
|---|---|---|
| Bin1-E17 | CCCTGAGAAAGGGAACAAGA | ATTCACAGTTGCGGAGAAGG |
| Bin1-E11 | TCAATGATGTCCTGGTCAGC | GCTCATGGTTCACTCTGATC |
| Dmpk | AACTACAGGAGGCCGAGGTC | TATAGGCCACCAACCCAGTG |
| Fhl1 | GAACGTGGAGTACAAGGGCA | TCGTGCCAGGATTGTCCTTC |
| Ldb3 | GCAAGACCCTGATGAAGAGG | GATGCTGGCAGTGGTTACG |
| Nrap | ACCGGCAGGACTTCCATAAG | CTGCCAACTTTGAGAGCGTG |
| Pdlim3 | GAAAAGCACACCCCTTCAAA | CCCGTCATTCACCAGATCCT |
| Pdlim7 | CCTGTTCAGAGCAAACCACA | TGAACTCTGTGCCCGTGA |
| Trim54 | CCTGCTCTCTCTGCAAGGTT | CTGCAACTTCTCCTCCTGCT |
| Aamdc | GGACAAATGAAAGTGCAAGGCT | AGACACCTCCTACCCTGACC |
| Dap3 | ATTGCCCCAGAGGAACTCTC | TTCTCATGTTGAAGCCAGTTG |
| Dnm1l | CTGCTTCTGCTGAGGCTGAT | TGCCTTTGGGACACTGTCTT |

| | | |
|---|---|---|
| Ppp3ca | CTGACACTGAAGGGCCTGAC | GAGGTGGCATCCTCTCGTTA |
| Ppp3cb ex10a | TTGGTTGCCCAATTTTATGG | CGGGCTGCAGCTGAACCTA |
| Ppp3cb ex13 | ACAGGGATGTTGCCTAGTGG | TCAGTGGTATGTGCGGTGTT |

**Table S3. Primer sequence used for qPCR**

| Gene | Forward (5'–3') | Reverse (5'–3') |
|---|---|---|
| Srsf2 | ACCTCCCTCAAGGTGGACAA | TTGTCGTGGAACCGGACG |
| Aurka | ATGACGCCACCCGAGTTTAT | GCTGTTCTCTGCTCGTCAAA |
| Fbxo5 | CAGCATCCCAACAAAACCTT | GCTCAGCCAGGATGTCTAGG |
| Nuf2 | TGGATTGCTTGCCTTCCTGT | TCTGGTCCTCCAAGTTCAAGC |
| Poc1a | CTGCTACCACCGTTGCCTTT | TGGTGGGACAGGCAAATCCA |
| Ranbp1 | AGATGAGTCCAACCACGACC | TTGACATCTCCAGTGCCTCG |
| Spdl1 | TAGACTCCCACCTCACAGGG | ATGTTGGCCGGACTATCCAC |
| Scn4b | GCTTCCCATGTACCTGTCGT | ATCCTGGATGTTTCGCTGTT |
| Stac2 | AGTCAGCGTTCACCTCTGGT | CAGTAGGTGGGGACTCCTTG |
| Synm | CTGGAGGATGAGAAGGAAGC | TGGCTACCTCCAAGCTGAGT |
| Tmod4 | CCTGAGACAACGTGACCAGA | CAGTGTAGGGCACCAAGTCA |
| Rplp0 | TAAAGACTGGAGACAAGGTGGGAG | AGAAAGCGAGAGTGCAGGGC |

**Table S4. SiRNA sequence used for RNA interference**

| | Sense | Antisense |
|---|---|---|
| siSRSF2 -#1 | CGAAGAUCCAAGUCCAAGUTT | ACUUGGACUUGGAUCUUCGTT |
| siSRSF2-#2 | UCCAGAUCAACCUCCAAGUTT | ACUUGGAGGUUGAUCUGGATT |
| siAurka-#1 | GAGCAGGUCAGAAGCCGGCACCAAA | UUUGGUGCCGGCUUCUGACCUGCUC |
| siAurka-#2 | CCAGAAACUUGGAGCAGGUCAGAAG | CUUCUGACCUGCUCCAAGUUUCUGG |
| siBin1-#1 | CAGCCAGUGAAGCAACCUCTT | GAGGUUGCUUCACUGGCUGTT |
| siBin1-#2 | CAGUGAAGCAACCUCCAGCTT | GCUGGAGGUUGCUUCACUGTT |
| siNC | UUCUCCGAACGUGUCACGUTT | ACGUGACACGUUCGGAGAATT |

**Table S5. Primer sequence used for ChIP-qPCR**

| Gene | Forward (5'–3') | Reverse (5'–3') |
|---|---|---|
| Aurka-ChIP | AAGGGACATGGCTGTTGAGG | ACAGAATGCAAACCCACTGAGA |
